# A prediction: granule cells can perform linearly non-separable computations

**DOI:** 10.1101/2021.04.02.438177

**Authors:** Romain D. Cazé

## Abstract

Multiple studies show how dendrites might extend some neurons’ computational capacity. Past works most often focus on pyramidal cells which have an extended dendritic tree where voltage can vary independently; pyramidal neurons’ dendrites emit local spikes turning dendritic branches into local non-linear subunits. However, these studies leave a large fraction of the nervous system unexplored. Here, we study a neuron with a modest dendritic and non-spiking dendrites. Granule cells’ dendrites do not spike and these cells’ membrane voltage remain constant over the neuron. This study uses a conjunction of Boolean algebra and biophysical modelling to predict that Granule cells can perform linearly non-separable computations. In a previous study, we found a linearly non-separable Boolean function possible to implement without dendritic spikes, we coined this computation the feature binding problem. In the present work, we use the intrinsic saturation of synaptic conductance to implement this computation in a biophysical model. An integrate and fire, cannot perform such type of computations. Confirming our prediction would change how we understand the nervous system.

## Introduction

We show here how dendrites can extend the computational capacity of all neurons, even the tiniest. We already knew that dendrites might extend the computational capacity of some pyramidal neurons; their extended dendrites capable of dendritic spikes changed the way we saw them (see [2] for one of the first article presenting this idea). Later suggested that we should model these neurons as a two layers network [7]. This theoretical model was further consolidated by experiments showing that we can see a pyramidal neuron as a collection of non-linear subunits [8]. Certain non-linearities can even allow a dendrite to implement the exclusive or (XOR) which is impossible for a point neuron [10]. A similar kind of nonlinearity, non-monotonic, was found in human pyramidal neurons [4]. But what about all other neurons with modest dendrites incapable of spiking?

Pyramidal neurons only represent a fraction of all neurons. The dendrites of cerebellar stellate cells cannot emit spikes, but they do saturate [1] and they can be decomposed into multiple independent subunits - with independent membrane voltages - turning them into two-stage units like the pyramidal neuron [9]. Previously we have shown that passive dendrites are sufficient to enable a neuron to perform linearly non-separable computations, for instance, the feature binding problem [3]. We focus here on granule cells which have on average four small (15*μ*m) dendrites and where membrane voltage remain equal everywhere [5]. Therefore dendrites and soma can be seen as a single unit contrary to what is happening in pyramidal neurons. In the present study, we demonstrate that these neurons can still implement a linearly non-separable computation. This opens the possibility to falsify the point neuron hypothesis with an experiment using existing techniques.

## Methods

### An integrate and fire with dendrites (the DIF)

We started from a leaky integrate and fire (LIF). This model has a membrane *V* modelled by the following equation:

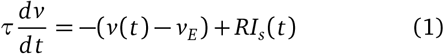

With *τ* = *20ms* is the neuron time constant, *v*(*t*) the membrane voltage at time *t* and *v_E_* = *–65mV* which sets the resting membrane voltage. *I_s_*(*t*) models the time varying synaptic inputs. It is described by

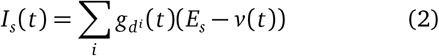

This current depends on the difference between *v*(*t*) the neuron voltage - equal everywhere - and *E_s_* the synaptic reversal potential (0*mV*) while *g_d^i^_* is the synaptic conductance in dendrite *i*. Each *g_d^i^_* is bounded between 0 and 1nS. Each *g_d^i^_* jumps up instantaneously to its maximal value for each incoming input spike and decays exponentially with time constant *τ_s_* = 1 ms. In a LIF all synaptic inputs are gathered into a single umbrella and *i* = 1. In the present study, we introduce the Dendrited Integrate and Fire (DIF) which includes at least two dendrites (*i* = 2). We group synaptic inputs into two groups each targeting a dendrite (one green and one blue, see Fig. 1). We used the Brian software to carry our simulations, the code is freely available on the git repository attached with this paper.

**Figure 1.**
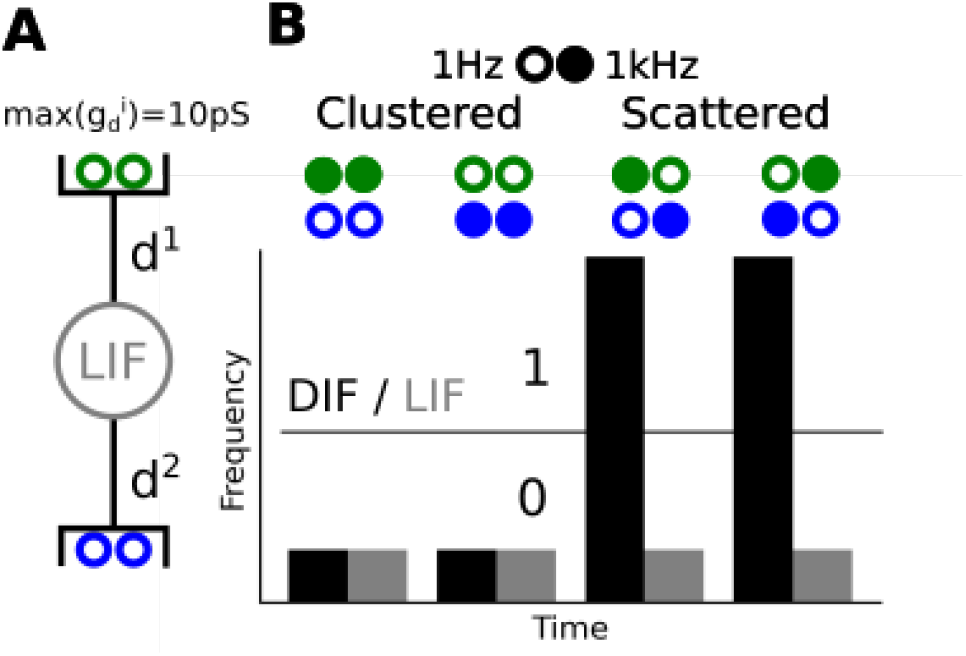
**A granule cell implementing a linearly non-separable computation. (A) A leaky integrate and fire (LIF) with two dendrites making it a dendrited integrate and fire (DIF), each half of the 4 synaptic inputs targets a distinct dendrite where** *g* **saturates at** 1*pS* **(B) Four stimulation scenarios, filled circles stand for a 1kHz input spike train while empty circles stand for 1Hz input spike train. Below, we plotted the response of the DIF (black) and a LIF (grey). We purposely removed the ticks label as the frequencies depend on the parameter of the model and input regularity. The parameters of the model can vary largely without affecting the observation. [6]**

### Boolean algebra reminder

First, let’s present Boolean functions:

#### Definition 1.

*A Boolean function of n variables is a function on* {0, 1}*^n^ into* {0, 1}, *where n is a positive integer*.

Importantly, we commonly assume that neurons can only implement linearly separable computations:

#### Definition 2.

*f is a linearly separable computation of n variables if and only if there exists at least one vector* 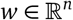 *and a threshold* 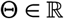 *such that:*

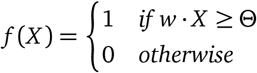

*where X* ∈ {0, 1}*^n^* is the vector notation for the Boolean input variables.

## Results

### The compact feature binding problem (cFBP)

In this section, we demonstrate a compact linearly non-separable computation that we are going to study. This computation is compact because it has four input/output lines. A LIF without dendrites cannot implement such computation.

The computation described in Tab. 1 is linearly inseparable. We call this computation the compact feature binding problem (cFBP).

**Table 1.**
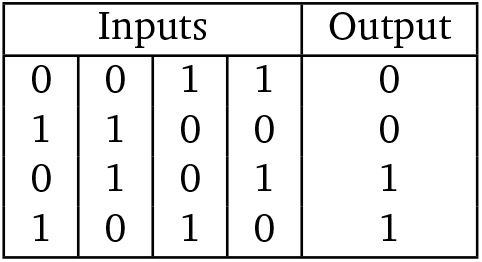
The truth table of a non-linearly separable computation

#### Proposition 1.

*The cFBP is linearly inseparable (non-separable)*

##### Proof.

The output must be 0 for two disjoint couples (1,2) and (3,4) of active inputs. It means that *w*_1_ + *w*_2_ ≤ Θ and *w*_3_ + *w*_4_ ≤ Θ we can add these two inequalities to obtain *w*_1_ + *w*_2_ + *w*_3_ + *w*_4_ ≤ 2Θ. However, the output must be 1 for two other couples made of the same active inputs (1,3) and (2,4). It means that *w_1_* + *w*_3_ > Θ and *w*_2_ + *w*_4_ > Θ we can add these two inequalities to obtain *w*_1_ + *w*_2_ + *w*_3_ + *w*_4_ > 2Θ. This yield a contradiction proving that no weights set exist solving this set of inequalities.

The cFBP is simple in two ways:

- Four input/output relations define this computation - the same number as the famous XOR (exclusive or).
- Contrary to the XOR it can be implemented with excitatory inputs and a monotone transfer function [3].

Notably, a neuron with a single dendrite can easily implement this computation. We proved how a ball-and-stick biophysical model can implement this computation in a previous study and we are going to use a similar method in the next section to implement the cFBP[3].

We can interpret the 0s and 1s in the truth table in at least two ways: (1) either the pre or post-synaptic neurons activates (2) or they reach a given spike frequency. In the following section, we will use the latter interpretation.

### Implementing the cFBP in a granule cell

We use local saturation in dendrites to implement the cFBP in a granule cell, the tiniest neurons in the brain. Their compactness makes them isopotential so we might wonder how local saturation can arise. Saturation has two possible origins: (1) a reduction in driving force can cause saturation like in [1], but (2) it can also be due to the intrinsic limitations in conductance. To better understand, imagine the neuron as a lake, the water level of the lake can depends on two things: (1) the level difference between the lake and rivers feeding into it, aka the driving force (2) the width of the river connected to the river. We are going to use this latter source of saturation to implement the cFBP in a granule cell.

We model a granule cell as a Dendrited Integrate and Fire (DIF) with two densrites as shown in Fig 1A. These cells receive high-frequency inputs from mossy fibres in the *kHz* range. Consequently, we consider a pre-synaptic input active when it fires a 1kHz regular spike-train and inactive if it fires at 1Hz. We stimulate our model in four different combination of inputs to reproduce the truth table from the previous section. You can observe on Fig 1B that when we bound *g*, it becomes possible to overcome granule cells compactness and to implement the cFBP. When we locally bound *g* the total current flowing into the neuron grows larger as we double the number of open gates if synaptic inputs distribute over the neuron. Having 2 independent *g* (*g*_*d*^1^_ and *g*_*d*^2^_) is key. If we have a unique *g_d_* the neuron remains a point and cannot identify inputs’ origins: it always fires at the same frequency (high or low depending on the value of the higher bound). The scenario with 2 dendrites differs when clustered inputs arrive on a dendrite the total input conductance *g* cannot exceed 1*nS*, in the scattered case the total input conductance can go up to *2nS* making the implementation of the cFBP possible.

## Conclusion/Discussion

In this brief report, we detailed a prediction: granule cells can implement linearly non-separable computations. These neurons have on average four dendrites, but we have shown here that two suffices. As we mentioned multiple times, multiple distinctly bounded *g* are crucial. We need a local saturation *g_d^i^_* to implement the cFBP

The experiment demonstrating this prediction seems straightforward. One would need to stimulate four distinct groups of mossy fibres following our different scenarios. We could then record how a group of granule cell respond using optogenetics reporting (i.e. calcium imaging). We predict that a significant part of Granule cells might implement the cFBP. This prediction could reveal the true potential of single neurons. The next step consists of looking at the network level as it was already done with spiking dendrites [**?**].

## Acknowledgements

I used “we” as science is a collective endeavour and this article came from interactions with A. Trah-Van-Min, T. Abrahamson, B. Gutkin and D. Digregorio. Discussions on a similar prediction had begun as early as 2013. I also want to acknowledge M. Humphries for his valuable comments on the early draft and Ms Marini-Audouard for the proof-reading before submission.

## Author contributions statement

R.C. wrote the initial draft, initiated the project, and made the initial simulations and figures.

